# The ion channel *ppk301* controls freshwater egg-laying in the mosquito *Aedes aegypti*

**DOI:** 10.1101/441592

**Authors:** Benjamin J. Matthews, Meg A. Younger, Leslie B. Vosshall

**Author notes:** denotes equal contribution (Benjamin J. Matthews, Meg A. Younger). Correspondence (L.B.V.).

## Abstract

*Aedes aegypti* mosquitoes are deadly vectors of arboviral pathogens including Zika, dengue, and yellow fever, and breed in containers of freshwater associated with human habitation^1,2^. Female *Ae. aegypti* lay eggs near freshwater because larval and pupal stages are aquatic^3^. They use volatile cues to locate water at a distance^4^, while at close-range they contact water to evaluate its suitability for egg-laying^4–7^. High salinity is lethal to mosquito offspring and therefore correctly laying eggs in freshwater is a crucial parenting decision made by female mosquitoes. Here we show that the DEG/ENaC channel^8–10^ *ppk301* is required for mosquitoes to exploit freshwater egg-laying substrates. When *ppk301* mutant females contact water, they do not lay eggs as readily as wild-type animals and are more likely to make aberrant decisions between freshwater and saltwater at concentrations that impair offspring survival. We used a CRISPR-Cas9-based genetic knock-in strategy combined with the Q-binary transactivator system^11^ to build genetic tools for labelling and imaging neurons in the mosquito. We found that *ppk301* is expressed in sensory neurons in legs and proboscis, appendages that directly contact water, and that *ppk301*-expressing neurons project to central taste centres. Using in vivo calcium imaging with the genetically-encoded calcium sensor GCaMP6s^12^, we found that *ppk301-*expressing cells respond to water but, unexpectedly, also to salt. This suggests that *ppk301* is instructive for egg-laying at low salt concentrations but that a *ppk301*-independent pathway is responsible for inhibiting egg-laying at high salt concentrations. Water is a key resource for insect survival and understanding how mosquitoes interact with water to control different behaviours is an opportunity to study the evolution of chemosensory systems. The new genetic tools described here will enable direct study of not only egg-laying, but also other behaviours in mosquitoes that influence disease transmission and enable comparative studies of insect biology more broadly.

---

*Ae. aegypti* females must take a blood-meal, typically from a human host, to develop eggs. Once proteins and other nutrients in the blood have been converted into eggs, she must find a body of freshwater suitable for laying her eggs. She first evaluates it using physical contact with all three pairs of legs and her proboscis. If a site is deemed suitable, she will lay eggs singly on the moist substrate above the waterline^5^ (Fig. 1a). When we used a mesh barrier to block access liquid water, females did not lay eggs, even though they had free access to the moist substrate (Fig. 1b,c). This suggests that direct contact with liquid water is necessary to stimulate egg-laying in *Ae. aegypti*.

**Figure 1.**
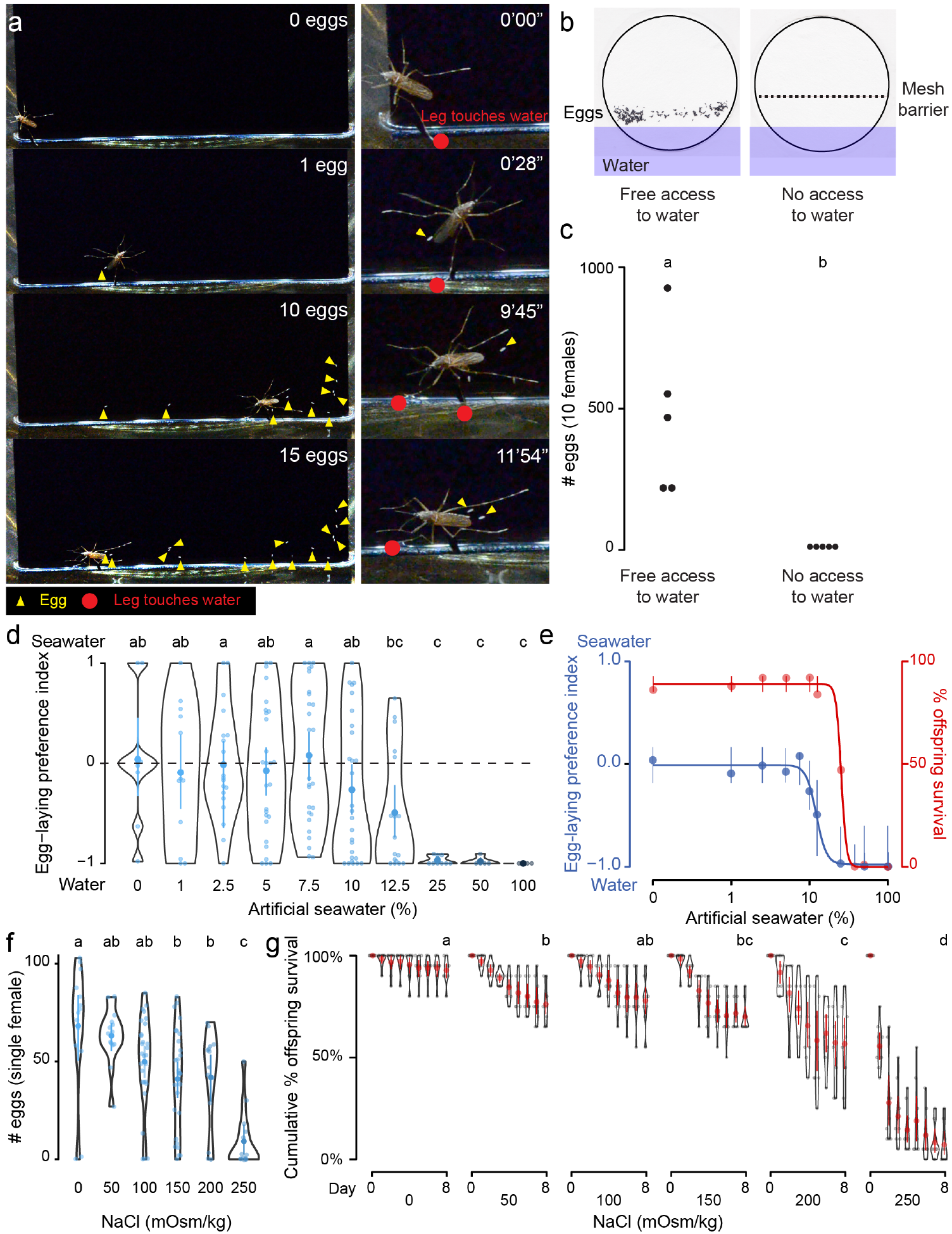
*Ae. aegypti* mosquitoes use freshwater contact to guide egg-laying, ensuring offspring survival. **a**, Still video frames of *Ae. aegypti* female with newly-laid eggs (yellow triangles) and contact between sensory appendages and liquid water (red circles) indicated. Right panels are magnification of animal in left panels. **b**, Egg-papers (black circles) from 10 females allowed to lay eggs for 18 hr with access to water (left) or access blocked by metal mesh (right). **c**, Eggs laid after 18 hr under conditions in **b**. n = 5; each dot represents data from a group of 10 females. **d**, Egg-laying preference of single females between water and increasing concentrations of artificial seawater. Mean ± 95% confidence interval. n = 4 – 29 / concentration. **e**, Four-parameter log-logistic curve fit to egg-laying preference in **d** (blue, n = 4 – 29 / concentration) or offspring surviving (red, n = 5 / concentration) in indicated concentrations of artificial seawater. Mean ± 95% confidence interval. **f**, Eggs laid by single females given access to liquid of the indicated NaCl concentration. Mean ± 95% confidence interval. n =14-28 / concentration. **g**, Cumulative survival of larvae in indicated concentration of NaCl. Mean ± 95% confidence interval. n=9 groups / concentration. In a given panel, data labelled with different letters are significantly different from each other; *P* < 0.05, two-way paired t-test (**c**) or ANOVA followed by Tukey’s HSD (**d**, **f**) or ANOVA followed by Tukey’s HSD of survival at day 8 (**g**).

As a consequence of their global spread, *Ae. aegypti* are faced with diverse habitats with a wide variety of potential egg-laying sites. For example, they can be found in abundance in a number of coastal regions rich with standing saltwater^13, 14^. To mimic the choice between freshwater and seawater in the lab, we developed a two-choice assay in which individual blood-fed females were placed in a container with a divided Petri dish with deionised water on one side and varying concentrations of a chemically-defined artificial seawater solution^15^ on the other (Fig. 1d). Mosquitoes showed no significant preference between deionised water and dilute seawater up to 10%, with individual mosquitoes either picking a solution at random or distributing their eggs between both solutions (Fig. 1d). However, they showed a strong dose-dependent aversion to higher concentrations of seawater, with an IC_50_ of 12.25% seawater (Fig. 1e). Females showed near-complete aversion to 25-100% seawater (Fig. 1d,e). These choices have consequences for the offspring. When we measured survival of offspring reared in varying concentrations of seawater, we found dose-dependent lethality (LD_50_ 25.23%) (Fig. 1e). To simplify the stimulus, we used sodium chloride (NaCl), the predominant salt in artificial seawater, in all subsequent experiments. Females showed dose-dependent inhibition of egg-laying with increasing concentrations of NaCl (Fig. 1f), suggesting that the preference for freshwater may be driven in part by an aversion to laying eggs in saltwater. Similar to artificial seawater, NaCl produced a dose-dependent decrease in offspring survival (Fig. 1g). This demonstrates that the female mosquito’s choice of freshwater or saltwater correlates with the survival of her offspring, making this an essential decision for the propagation and fitness of the species.

In a search for genes that control *Ae. aegypti* freshwater egg-laying, we reasoned that animals carrying a mutation in an egg-laying preference gene would fail to lay eggs in freshwater, inappropriately lay in saltwater, or both. *pickpocket* (*ppk*) genes encode cation channel subunits that function as putative mechanoreceptors and chemoreceptors for a wide array of stimuli including pheromones, liquid osmolality, and salt^8–10^. In a previous study, we generated CRISPR-Cas9 mutations in four *ppk* genes^16^ (Extended Data Fig. 1) because we considered these as possible candidates for controlling egg-laying. We first measured the ability of these mutant strains to blood-feed, and found that all four mutants were attracted to and engorged fully on the blood of a live human host (Fig. 2a).

To test egg-laying behaviour, we introduced single blood-fed females of each strain into egg-laying vials containing a small amount of water and a filter paper as an egg-laying substrate. One mutant, *ppk301*, an orthologue of the *Drosophila melanogaster ppk28* low-osmolality sensor^17, 18^, showed a defect in egg-laying. Fewer than 40% of *ppk301* mutants laid more than 10 eggs (Fig. 2b). To exclude the possibility that this defect was due to an inability to convert blood into developed embryos, we counted the number of mature eggs in ovaries and confirmed that there was no difference between wild-type and *ppk301* mutants (Fig. 2c).

To investigate freshwater egg-laying preference in the *ppk* mutants, we introduced single female mosquitoes into individual chambers containing a choice between freshwater (0 mOsm/kg NaCl) or 200 mOsm/kg NaCl. Even when *ppk301* mutant animals laid eggs, they laid fewer eggs than wild-type on water and more eggs than wild-type on 200 mOsm/kg NaCl (Fig. 2d). We assayed these four *ppk* mutant strains across a range of NaCl concentrations and found a significant reduction in aversion to salt solution only in the *ppk301* mutant, as measured by the proportion of animals laying eggs primarily on salt solution (Fig. 2e,f). Together, these data suggest that mutations in *ppk301* disrupt freshwater egg-laying in two distinct ways: by dramatically reducing the drive to lay eggs in suitable, low-salt, substrates and reducing the aversion to concentrations of NaCl that are lethal to their offspring.

If *ppk301* mutant animals fail to sense water, which normally triggers mosquitoes to lay an entire clutch of eggs in a short timespan, we hypothesized that these mutants would slowly lay eggs when housed in close proximity to water for many days. To address whether *ppk301* mutant animals will lay eggs on water given sufficient time, we introduced blood-fed females into egg-laying vials containing water and monitored the number of eggs laid per female per day over seven days. While the vast majority of wild-type and heterozygous animals laid all of their eggs within the first two days of being introduced into egg-laying vials, *ppk301* mutant animals did not. Instead, the mutants showed increased variability in the time of egg-laying initiation and a tendency to spread egg-laying out over many days (Fig. 2g-j). This slow, sustained, and variable egg-laying behaviour on freshwater is consistent with defect in sensing the water that triggers egg-laying.

**Figure 2.**
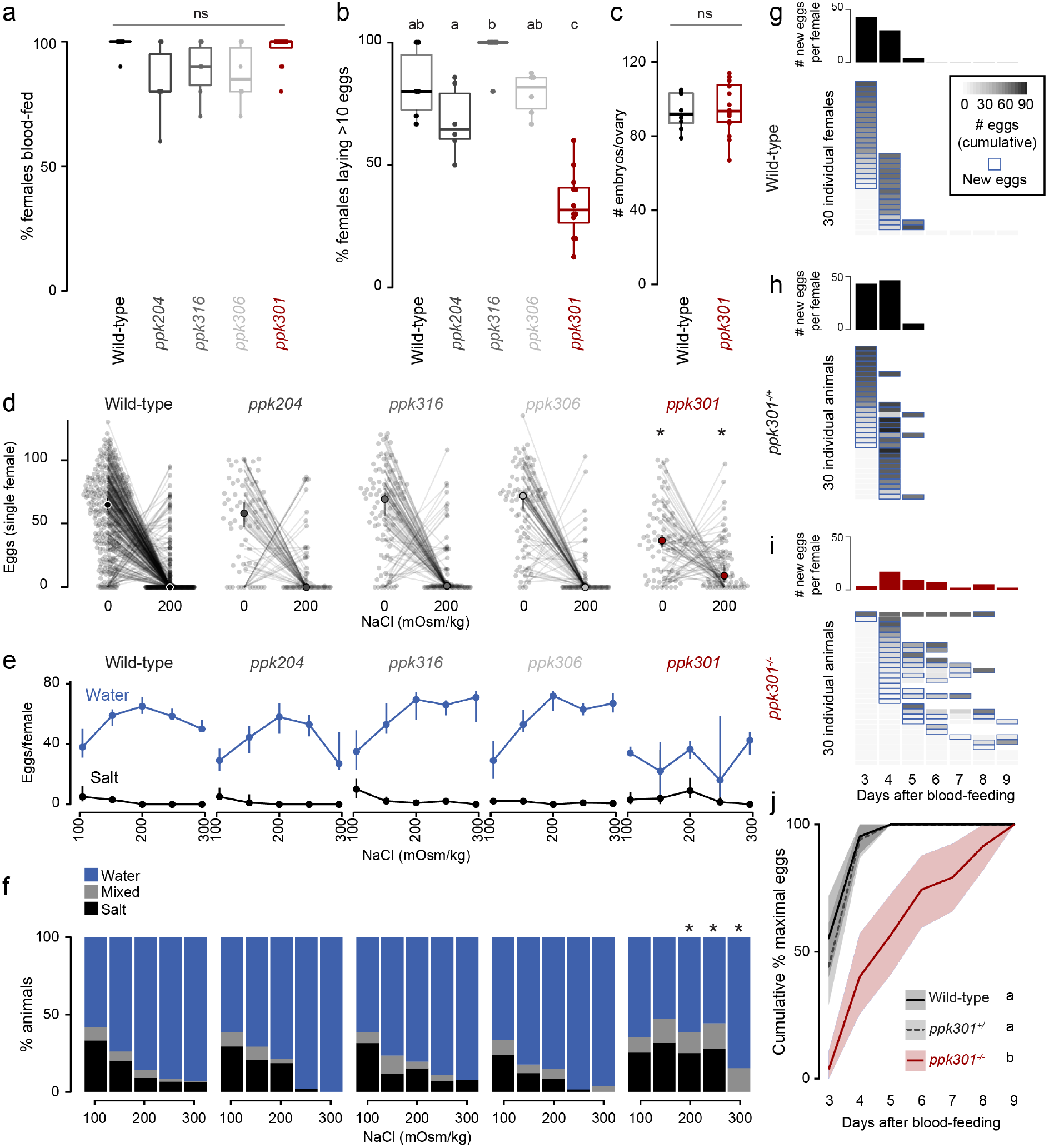
Mutations in the *ppk301* ion channel disrupt egg-laying preference and timing. **a, b** Females of indicated genotype who blood-fed (**a**) or laid > 10 eggs given 18 hr to lay (**b**). n = 6 - 12 groups / genotype. **c**, Embryos / ovary 72 hr post-blood-meal. n = 8 - 16 / genotype. **d-f**, Eggs laid by single females of the indicated genotype in the indicated NaCl concentration. Only females laying > 10 eggs are presented. In **d** lines connect data from individual animals (dots). Median (points) ± 95% confidence interval (bars). n = 65 - 314 animals / genotype (**d**) or n = 3 - 314 animals / genotype (**e**) per concentration. **f**, proportion of animals binned into three behavioural groups by eggs laid on each substrate. **g-i**, Eggs laid in 0 mOsm/kg NaCl by single females of the indicated genotype. n = 30 females/genotype. Blue outline indicates days on which new eggs were laid and histograms indicate mean new eggs (per female). **j**, Summary of data in **g-i**. Mean (line) ± 95% confidence interval (shaded bars). In **b**, data labelled with different letters are significantly different from each other *P* < 0.05, in * indicate difference from wild-type *P* < 0.05 (**d**) P < 0.01 (**f**); ns = not significant, ANOVA followed by Tukey’s HSD (**a, b, e, h**), unpaired t-test (**c**), and chi-squared test corrected for FDR (**f**). Boxplots in **a-c** indicate median and 1st and 3rd quartiles, whiskers extend to 1.5x interquartile range.

If *ppk301* directly senses the osmolality or salinity of liquid, we would expect it to be expressed in the sensory appendages that contact water. At the inception of this project, genetic tools for labelling, monitoring, and manipulating neurons in *Ae. aegypti* did not exist. To address this gap, we developed new genetic tools in the mosquito to label all *ppk301*-expressing neurons, and to image neuronal activity at sensory neuron terminals using the genetically-encoded calcium sensor GCaMP6s^12^. We adapted an approach in which a T2A ‘ribosomal skipping’ peptide is used to express multiple independent protein products from a single RNA transcript^19, 20^. We first tested the efficiency of T2A in *Ae. aegypti* by generating a transgene containing a membrane-targeted mCD8:GFP fusion protein and a nuclear-targeted dsRed:NLS fusion protein separated by T2A and driven from the *Ae. aegypti* polyubiquitin promoter^21^ (Fig. 3a). Confocal microscopy revealed complete subcellular separation of the two fluorophores in individual larval body-wall cells (Fig. 3a). This demonstrates that T2A functions efficiently to prevent peptide bond formation in *Ae. aegypti* and can be used as a tool to independently express multiple gene products from a single locus.

To build a flexible system for expressing a diverse array of effector transgenes, we employed the Q-binary expression system for transgene amplification^11^, which has been successfully implemented in *Anopheles gambiae* malaria mosquitoes^22^. We used CRISPR-Cas9 with the same guide RNA as the *ppk301* mutant to introduce an inframe T2A sequence into the *ppk301* locus followed by the QF2 transcriptional activator (Fig. 3b). This is predicted to cause a loss-of-function mutation in *ppk301* and simultaneously express QF2 in all *ppk301*-expressing cells.

We also generated two transgenic QUAS effector strains. The first is a QUAS response element driving the expression of both cytosolic tdTomato and GCaMP6s (*15xQUAS-tdTomato-T2A-GCaMP6s*), which allows us to simultaneously label neurons and image their activity. The second drives expression of membrane-bound GFP (*15x-QUAS-CD8:GFP*), allowing us to reveal the complete morphology of the neurons in which it is expressed (Fig. 3c,i). We looked for expression of the *ppk301>tdTomato-T2A-GCaMP6s* reporter in the appendages that contact water during egg-laying (Fig. 3d) and found expression in sensory neurons innervating trichoid sensilla in the labellar lobes of the proboscis (Fig. 3e) and in the legs, primarily in the distal tarsal segments (Fig. 3f).

The mosquito central nervous system consists of a brain and a ventral nerve cord^23–25^ (Fig. 3g). To facilitate our understanding of the neuroanatomy of *Ae. aegypti* in general, and the projection pattern of the *ppk301* driver line in particular, we built a threedimensional female mosquito brain atlas, in which we identified and named the major neuropils in accordance with nomenclature established by the Insect Brain Name Working Group^23^ (Fig. 3h and http://mosquitobrains.org). Projections of *ppk301*-expressing sensory neurons in the head extend processes specifically to the subesophageal zone (Fig. 3h,j). Two nerves enter each hemisphere of the subesophageal zone, arising from the proboscis and the pharynx (Fig. 3j). Labelling was absent in genetic controls expressing either QF2 driver or QUAS effector alone (Extended Data Fig. 2d-k). Additionally, each leg sends projections into the ventral nerve cord, with nerves running into each neuromere (Fig. 3k). Both brain and ventral nerve cord innervation patterns are consistent with these neurons mediating taste sensation^26^. We noted that projections of *ppk301*-expressing neurons are also present in the male brain and ventral nerve cord (Extended Data Fig. 2b,c), consistent with a role for *ppk301* in behaviours other than egg-laying.

We hypothesized that if *ppk301*-expressing neurons were promoting egg-laying, they should be maximally activated by water, and we set out to test this by functional calcium imaging in response to multiple behaviourally relevant NaCl solutions within one animal. We developed an in vivo calcium imaging preparation with GCaMP6s^12^ in ventral nerve cord sensory afferents of a mosquito presented with water or NaCl solutions on a single foreleg (Fig. 4a,b). This appendage was chosen because it most frequently contacts water during egg-laying (Fig. 1a). We imaged the prothoracic segment of the ventral nerve cord with two-photon microscopy, using a custom fluidics device to deliver and retract liquids to the foreleg. *ppk301*-expressing neuronal projections in the ventral nerve cord were identified by tdTomato expression (Fig. 4c), which was also used to determine the region of interest for calcium imaging analysis.

**Figure 3.**
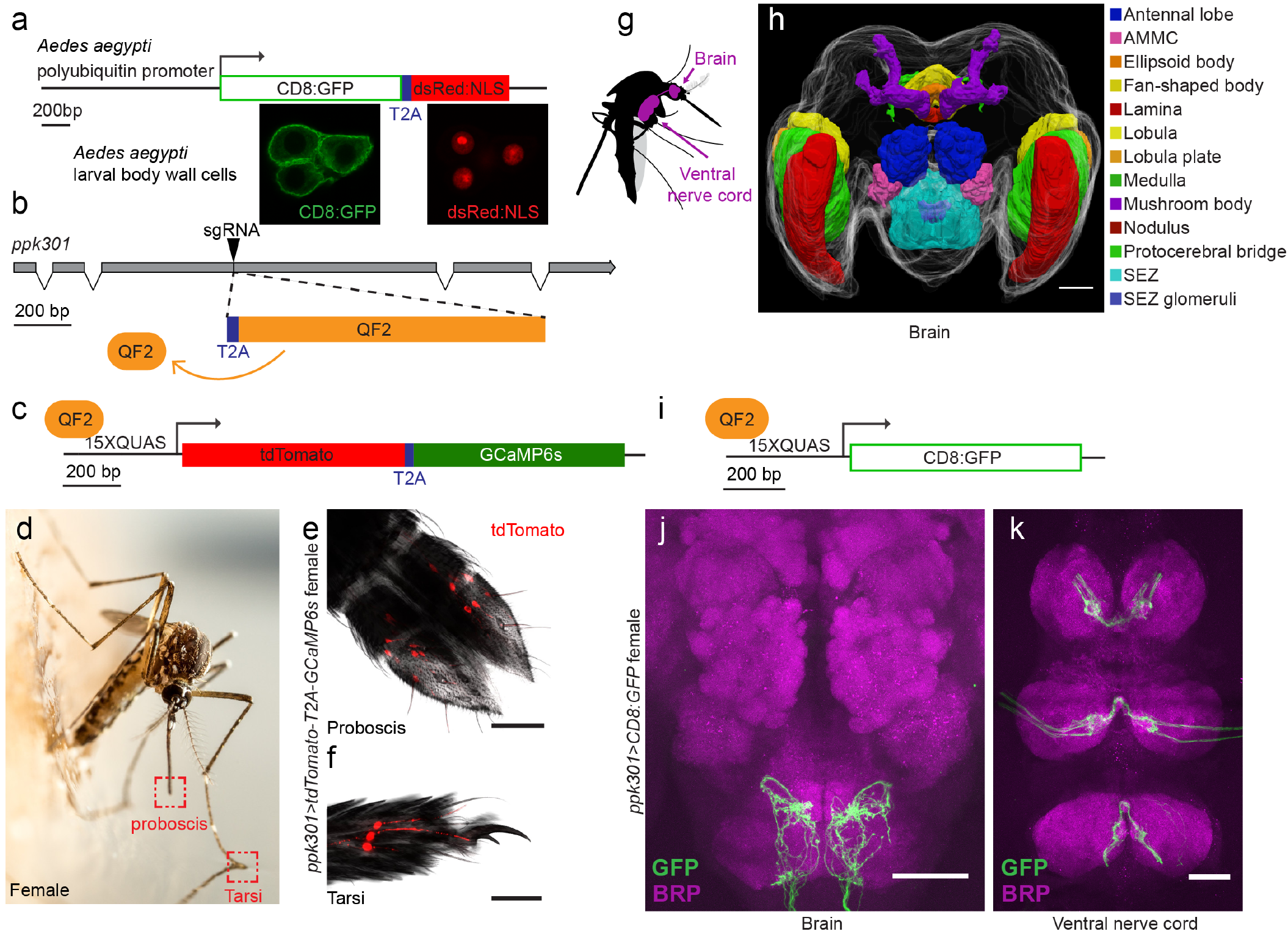
In-frame insertion of T2A-QF2 into the *ppk301* locus labels sensory neurons that project to central taste centres. **a**, Top, construct used to test the efficacy of T2A in *Ae. aegypti*. Bottom, confocal images of 3 *Ae. aegypti* body wall cells expressing both CD8:GFP (green) and dsRed:NLS (red). **b**, Diagram of coding region of *ppk301* locus with exons (grey boxes), introns (connecting lines), and CRISPR-Cas9 gRNA site used to insert T2A (blue) and QF2 (orange). **c**, Map of *QUAS-tdTomato-T2A-GCaMP6s* transgene. **d**, Female mosquito laying an egg on a moist substrate, highlighting proboscis and tarsal appendages that contact water (red boxes). Photo: Alex Wild. **e**, **f**, Confocal image tdTomato expression in *ppk301>tdTomato-T2A-GCaMP6s* proboscis (**e**) and tarsi (**f**) with transmitted light overlay. **g**, Cartoon of mosquito adult neural tissues. **h**, 3D-reconstruction of female *Ae. aegypti* brain. Different regions of the brain are identified by Bruchpilot (Brp) immunostaining and homology to other insects. AMMC = Antennal Mechanosensory and Motor Centre, SEZ = subesophageal zone. i. Map of *QUAS-CD8:GFP* transgene **j**, **k**, Expression of *ppk301>CD8:GFP* in female brain (**j**) and ventral nerve cord (**k**) Scale bars: 50 μm.

We observed low GCaMP6s fluorescence at baseline and an increase in fluorescence in every trial where water was presented (Fig. 4c-e), with no apparent desensitization across trials (data not shown). Only the ipsilateral side of the ventral nerve cord innervated by the stimulated leg showed activation (data not shown). In *D. melanogaster*, cells that express the *ppk301* orthologue, *ppk28*, respond to water but are inhibited by high osmolality solutions including NaCl^17,18^. The population response in *ppk301*-expressing afferents in the mosquito is functionally different, showing robust responses to water and activation by salt solutions (Fig. 4c-e), with the strongest response at 200mOsm/kg NaCl. This suggests that mosquito *ppk301*-expressing cells are either multi-modal^27^ and respond to both water and salt or are functionally heterogeneous with distinct neurons responding to each cue.

The observation that *ppk301*-expressing cells are activated by both water and high salt is intriguing because wild-type females avoid laying eggs in high salt. To understand how female mosquitoes interact with these different stimuli, we monitored real-time behaviour of individual females offered either water or 300 mOsm/kg NaCl over 40 min by scoring their contact with liquid and individual egg-laying events. Both wild-type and *ppk301* mutant mosquitoes contacted water, but only wild-type females consistently transformed these touches into egg-laying events (Fig. 4f). When offered 300 mOsm/kg salt solution, both wild-type and *ppk301* mutant mosquitoes touched liquid but neither genotype reliably laid eggs (Fig. 4f). These results are consistent with a model (Fig. 4g) in which activation of *ppk301* cells by water gates rapid and reliable egg-laying. Animals lacking *ppk301* fail to detect the water activation signal and show delayed and intermittent egg-laying. This model predicts that *ppk301*-expressing cells gate egg-laying at low NaCl concentrations, but as NaCl concentrations increase, egg-laying is inhibited by an independent noxious salt-sensing pathway that overrides water activation of *ppk301*-expressing cells (Fig. 4g). We predict mutations that disrupt this noxious salt sensor would yield mosquitoes that show indiscriminate egg-laying on a high-salt substrate.

We have identified a mutation disrupting freshwater egg-laying behaviour in a mosquito, providing an entry point into the neural circuitry underlying the most important parenting decision a female mosquito ever makes. In contrast to *ppk301*-expressing cells in the mosquito ventral nerve cord, which are activated by both water and high salt, neurons in the *D. melanogaster* proboscis expressing the orthologous *ppk28* gene are activated by water but inhibited by dissolved solutes^17, 18^. It will be interesting to discover if *Ae. aegypti* ppk301 responds both to water and salt, or whether *ppk301* cells express additional receptor(s) that respond to salt. DEG/ENaC channels are trimers^28^, and it remains to be seen whether ppk301 is part of a homomeric channel that is tuned to water and NaCl or if the ppk301 subunit contributes to multiple receptors with different properties. It is also unknown whether the population of *ppk301*-expressing cells in the legs and proboscis are functionally heterogeneous as has recently been shown in *Drosophila*^29^.

Some strains of *Ae. aegypti* have begun to exploit brackish water for egg-laying^13, 14^, and our work provides an opportunity to understand how evolution may act on genes and circuits to shift adult behavioral preference and larval salt tolerance to exploit a novel ecological niche. The tools described here make it possible to visualize neuronal anatomy and activity within molecularly-identified cell types and identify the genetic and neural circuit substrates of many mosquito behaviours. The development of similar tools and reagents in other insect species of public health, agricultural, or ethological interest will broaden our view of insect biology and facilitate comparative studies of the genes and circuits underlying evolutionary adaptations in insects. These behaviours contribute to the spread of deadly pathogens and understanding the underlying biology of these behaviours will contribute to control efforts, in addition to providing an invaluable comparative window into the organization and function of insect nervous systems.

**Figure 4.**
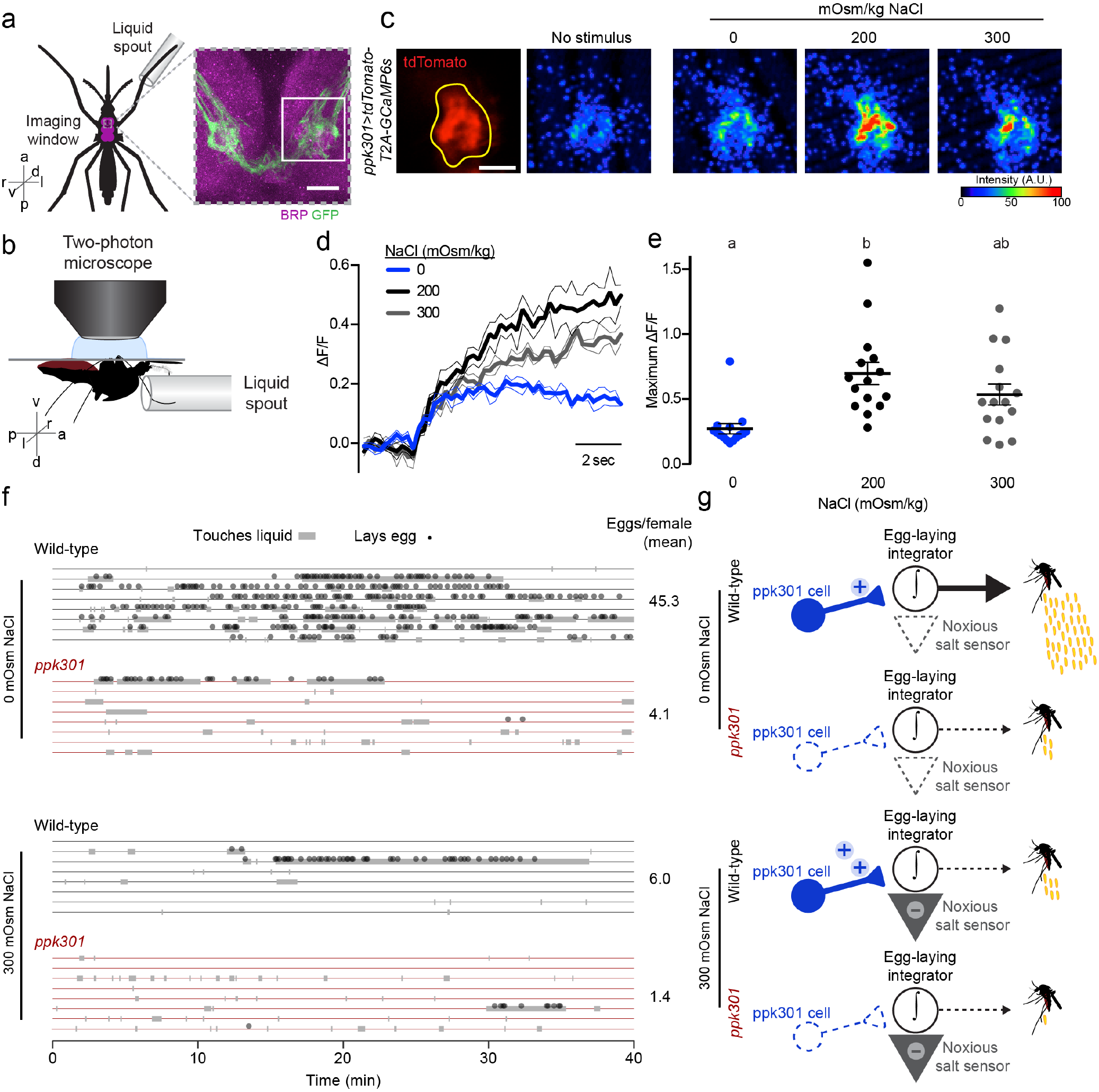
*ppk301* neurons respond to water and salt, pointing to integration of water and salt cues in driving egg-laying preference. **a**, Left, ventral view schematic of mosquito during imaging showing ventral nerve cord (magenta) and imaging window (grey box). Right, confocal image of the approximate area contained in the imaging window. White box highlights the approximate area in **c**. Image from *ppk301>CD8:GFP* animal. **b**, Side view schematic of ventral nerve cord imaging setup. **c**, Left, two-photon image of tdTomato (red) and region of interest (yellow). Right: Representative GCaMP6s fluorescence increase to indicated NaCl concentrations compared to baseline (scale: arbitrary units). **d**, GCaMP6s traces from an animal presented with 3 concentrations of NaCl until end of trial, 3 sweeps each. Mean (thick lines) ± S.E.M. (thin lines). **e**, Maximum ΔF/F (arbitrary units) for individual sweeps. n = 15 sweeps / condition from n = 5 animals. Data labelled with different letters are significantly different from each other; *P* < 0.05, one-way ANOVA with Tukey’s HSD. **f**, Raster plot of eggs laid (circles) and physical contact with liquid (grey boxes) by 8 individual females of the indicated genotype over 40 min observation period in the indicated NaCl solution. **g**, Model for *ppk301* control of freshwater egg-laying. Scale bars: 20μm (**a**) or 10μm (**b**).

## SUPPLEMENTAL INFORMATION

All raw data in the paper are provided in Supplemental Data File.

## ACKNOWLEDGMENTS

We thank Raphael Cohn, Emily J. Dennis, Andreas Keller, Kevin J. Lee, Gaby Maimon, Lindy McBride, Vanessa Ruta, Vikram Vijayan, Nilay Yapici, and members of the Vosshall Lab for comments on the manuscript. Funding for this study was provided by Jane Coffin Childs Postdoctoral Fellowships (B.J.M and M.A.Y.), The Grass Foundation (M.A.Y.), a Leon Levy Neuroscience Fellowship (M.A.Y.), and by a pilot grant and postdoctoral fellowship from the Kavli Institute for Systems Neuroscience (M.A.Y.). We thank the following research assistants, and high school, college, and rotation PhD students for assistance in collecting egg-laying data: Julia Canick, Emily Dennis, Solomon Dworkin, Katie Kistler, Stephanie Marcus, Nicholas Urban Schwartz, Russell Shephard Eva Shrestha, Krithika Venkataraman, Josh Zeng. We thank Alison Ehrlich and Zachary Gilbert for assistance with molecular biology and mosquito husbandry; Gloria Gordon and Libby Mejia for expert mosquito rearing; Victoria Danan, Nick Didkovsky, Nicolas Renier for contributions to the mosquitobrains.org project; Raphael Cohn, Gaby Maimon, Vanessa Ruta, John Tuthill, and Ari Zolin for guidance on calcium imaging; James Petrillo, Daniel Gross, Peer Strogies of the Rockefeller Precision Instrumentation Technology core for design and fabrication of egg-laying assays and a device for calcium imaging; Rob A. Harrell II at the Insect Transgenesis Facility at the University of Maryland for CRISPR-Cas9 and transgene injections. We thank Olena Riabinina and Christopher Potter for generously making unpublished QF2/QUAS reagents, data, and protocols available to us prior to publication. L.B.V. is an investigator of the Howard Hughes Medical Institute.

## AUTHOR CONTRIBUTIONS

B.J.M., M.A.Y., and L.B.V. conceived the project, wrote the manuscript and produced the figures. B.J.M. carried out experiments in Fig. 1-2, Fig. 3e,f, Fig. 4e,f, Extended Data Fig. 1; M.A.Y. carried out experiments in Fig. 3a, g-k, Fig. 4a-e, and Extended Data Fig. 2. B.J.M. generated ppk301-T2A-QF2 Fig. 3b. B.J.M. and M.A.Y. generated all transgenic reagents Fig. 3a,c,i.

## MATERIALS AND METHODS

### Contact for reagent and resource sharing

Further information and requests for reagents should be directed to and will be fulfilled by the lead contact, Leslie Vosshall.

### Mosquito rearing and maintenance

*Aedes aegypti* wild-type laboratory strains (Liverpool; LVP) were maintained and reared at 25 - 28°C, 70-80% relative humidity with a photoperiod of 14 hr light: 10 hr dark as previously described^30^. Adult mosquitoes were provided constant access to 10% sucrose. Adult females of all genotypes were blood-fed on a single human subject during mutant generation, stock maintenance, and for behavioral assays. Blood-feeding procedures with live hosts were approved and monitored by The Rockefeller University Institutional Animal Care and Use Committee (IACUC protocol 15772) and Institutional Review Board, (IRB protocol LV-0652). Human subjects gave their written informed consent to participate.

### Blood-feeding for egg-laying behaviour

All *Ae. aegypti* mosquitoes used in egg-laying assays were housed in mixed-sex cages and were between 7-14 days old. Female mosquitoes were blood-fed by giving them direct access to a human limb introduced in or on the mesh wall of a cage. Fully engorged female mosquitoes were selected within 24 hr of blood-feeding and housed in insectary conditions with ad libitum access to 10% sucrose before being introduced into behaviour assays. Unless otherwise indicated, assays were performed beginning 96 hr post-blood-meal and continuing for ~18 hr.

### Mesh barrier assay

10 blood-fed female mosquitoes were introduced by mouth aspiration into a standard BugDorm rearing cage (30 cm^3^) containing a 10% sucrose wick. Two egg-laying cups were present in each cage, each containing deionised water and a 55 mm-diameter partially submerged Whatman filter paper (Grade 1; 1001-055). In one of the cups, a metal mesh barrier placed ~1 cm above the water line prevented direct access of the mosquitoes to liquid, although the filter paper remained partially submerged and moist. Mosquitoes were allowed to lay eggs overnight (18 hr), after which the filter papers were dried and eggs counted.

### Seawater preference assay

Artificial seawater was mixed according to the recipe of Kester et al., 1967^15^ with lab-grade chemicals from Sigma-Aldrich. Specific dilutions were made, by volume, with deionised water. Single blood-fed females were introduced into a chamber comprised of a length of transparent acrylic tubing (inner diameter 9.525 cm) cut to 12 cm in height with a wire mesh grid glued to one end as a ceiling containing a two-sector 90 mm divided Petri dish with 10 mL of deionised water on one side and 10 mL of a specific dilution of artificial seawater on the other. As an egg-laying substrate, a 1 cm tall strip of seed germination paper (Anchor Paper; SD7615L) was wrapped around the outer diameter of each half of the Petri dish, partially submerged in the liquid. Animals were introduced by a mouth aspirator through a hole in the top of the chamber, which was then plugged with a cotton ball. Containers were stored under insectary conditions and mosquitoes were allowed to lay eggs overnight (~18 hr), after which the paper strips were dried and eggs counted.

### Seawater survival assay

Wild-type mosquitoes were hatched in ‘hatch broth’ (deoxygenated water containing ground Tetramin fish food). Approximately 1 day after hatching, 20 larvae were transferred into a small plastic cup containing 25 mL of a specific dilution of artificial seawater. Cups were examined each day, dead larvae removed, and ground Tetramin pellets added for food as needed. Animals successfully completing the transition to pupal stage by 8 days post-hatch were scored as surviving offspring.

### Single-female modular egg-laying assay

A multi-animal egg-laying assay was created out of sheet acrylic, comprising modular trays each with 14 single-animal chambers. Each chamber comprised two angled 50 mm Petri dishes each containing 2 mL of liquid and a 47 mm diameter filter paper (Whatman, Grade 1 Qualitative Filter Paper).

We next developed a standardized imaging setup to automatically count eggs from each Petri dish. Each dish was placed onto a LED light panel (SuperbrightLEDs.com item #2020) and images were captured with a Raspberry Pi and associated camera. Images were thresholded and pixels counted for each dish of each chamber. The number of eggs corresponding to each image was determined by dividing the average pixel value of eggs determined from a manually-counted set of 20 test images. The concordance between manual counting and automatic pixel-based counting was r^2^ = 0.96. A parts list and schematics of the egg-laying chambers and the image acquisition and thresholding code can be found on http://github.com/VosshallLab/MatthewsYoungerVosshall2018/.

For studies of the effect of NaCl on egg production (Fig. 1f), each dish was filled with the same concentration of NaCl. The osmolality of each solution was measured using a Wescor model 5520 vapour pressure osmometer. Blood-fed female mosquitoes were cold anesthetized and single animals introduced into each chamber by mouth aspiration. Animals were allowed to lay for 18 hr and dishes were imaged to calculate egg numbers. For two-choice assays (Fig. 2d-f), assays were performed identically, except that the two dishes contained deionised water and a NaCl solution of a specific osmolality. The position of each solution was varied for each chamber. Experiments were performed blind to genotype and data included only for those animals who laid more than 10 eggs.

### NaCl survival assay

Wild-type mosquitoes were hatched in hatch broth. Approximately 1 day after hatching, 20 larvae were transferred into a small plastic cup (VWR HDPE Multipurpose Containers; H9009-662) containing 25 mL of a specific concentration of NaCl, prepared as above. Cups were examined each day, dead larvae removed, and ground Tetramin pellets added for food as needed. The number of pupae and larvae remaining alive were counted each day, and cumulative survival was calculated for these offspring.

### Egg-laying timing: vial assay

To measure the timing of egg-laying across days, individual blood-fed mosquitoes were introduced into egg-laying vials 48 hr after a blood-meal. Mosquitoes were transferred to a fresh vial every 24 hr and eggs from the previous day were counted and recorded. Experiments were performed blind to genotype.

### Egg-laying timing: culture flask assay

To measure the timing of egg-laying and liquid touching, single animals were introduced into a 50 mL cell culture flask containing 10 mL of either deionised water or 300 mOsm/kg NaCl, and a 1” × 2” strip of seed germination paper, partially submerged. Animals were video recorded for 40 min using a Nikon D7000 SLR with a macro lens. Four flasks were recorded simultaneously, with each set containing 1 replicate of the following conditions: wild-type, 0 mOsm/kg NaCl; wild-type, 300 mOsm/kg; *ppk301*, 0 mOsm/kg; *ppk301*, 300 mOsm/kg. Videos were manually scored, blind to genotype and condition, for physical contact with liquid and the appearance of freshly laid eggs. Data on physical touches were recorded in 5 s intervals, with each interval scored as ‘touch’ if a single frame revealed physical contact between any appendage of the mosquito and the liquid. Videos were scored blind to genotype and condition.

### Curation and expression estimates of ppk gene family in *Ae. aegypti*

To identify members of the ppk gene family in *Ae. aegypti* and *An. gambiae*, we performed two complementary analyses: 1) using *D. melanogaster* ppk sequences as queries, we performed BLASTp against all translated protein-coding genes identified in AaegL5 (GCF_002204515.2_AaegL5.0_protein.faa) and 2) ran interproscan v5.29.68.0^31^ against the same database of translated protein-coding genes. We took all genes that were reciprocal best hits with the *D. melanogaster* ppk family via blastp and were annotated by interproscan as ‘Amiloride-sensitive sodium channel.’ We repeated this analysis for the *An. gambiae*, PEST strain geneset downloaded from Vectorbase (Anopheles-gambiae-PEST_PEPTIDES_AgamP4.9.fa). We re-named *Ae. aegypti ppk* genes by giving them a 3-digit identifier corresponding to their chromosomal position. The first digit represents the chromosome (1, 2, or 3), while the next two digits represent its relative position on that chromosome from the left (p) arm to the right (q) arm according to coordinates found on NCBI.

We next built a phylogenetic tree to visualize the relationship between these genes across these three species, we selected the longest single isoform for genes predicted to encode multiple protein-coding isoforms and performed multiple sequence alignment using clustal-omega v1.2.3^32^, including a human ASIC channel and *Caenorhabditis elegans* MEC4 sequence for comparison. A maximum-likelihood-estimate phylogenetic tree was constructed using PhyML v3.0^33^ with default parameters and 100 bootstrap iterations, and manually re-rooted on MEC4 for presentation. A table of all genes and sequences incorporated in this analysis, with previous accessions, is presented in the Supplementary Data File.

### Transcript abundance estimates of *Ae. aegypti ppk* genes

To visualize transcript abundance of each predicted *ppk* gene across tissues, we utilized published data^34, 35^. Heatmaps were generated as described^35^ and presented as log10(TPM + 1) of the mean expression value for all replicates of the indicated tissue using the heatmap.2 function of the gplots v3.0.1^36^ package in R v3.5.0^37^.

### Generation of mutant and transgenic mosquito strains

*ppk* loss-of-function alleles were generated and described previously^16^. All new strains generated in this paper were injected at the Insect Transformation Facility at the University of Maryland.

<u>PUb-mCD8:GFP-T2A-dsRed:NLS-SV40</u> was generated by Gibson assembly, using the following fragments: Plasmid backbone and MOS arms from a standard transformation vector (*pMos-3xP3-dsRed*); *dsRed* open reading frame amplified from the same vector by polymerase chain reaction (PCR); *mCD8-GFP* open reading frame PCR-amplified from a synthesized vector (Genscript); *Ae. aegypti PUb* promoter PCR-amplified from *PSL1180-HR-PUb-ECFP* (Addgene #47917). T2A and NLS sequences were added through PCR. 1000 *Ae. aegypti* embryos (LVP strain) were injected with the plasmids and a Mos helper plasmid. Two stable lines were recovered with qualitatively similar expression patterns.

<u>*ppk301-T2A-QF2*</u> We initially attempted to use constructs with gene-specific promoter fragments to drive transgenic constructs, including the broadly expressed polyubitiquin promoter^21^, but these either did not express at all or were expressed sporadically and mostly in non-neuronal cells^38^. We also had no success with the Gal4/UAS^39^ system in *Ae. aegypti*. We therefore developed techniques to knock in QF2 into the *ppk301* locus.

*ppk301-T2A-QF2* was generated with a sgRNA targeting exon 2 of the *ppk301* locus (*ppk301-sgRNA-1*, target sequence with PAM underlined: GGTTGGCAGTTGAGTCC**<u>CGG</u>**). sgRNA DNA template was prepared by annealing oligonucleotides as described^16^. In vitro transcription was performed using HiScribe Quick T7 kit (NEB, E2050S) following the manufacturer’s directions and incubating for 2 hr at 27°C. Following transcription and DNAse treatment for 15 min at 37°C, sgRNA was purified using RNAse-free SPRI beads (Ampure RNAclean, Beckman-Coulter A63987), and eluted in Ultrapure water (Invitrogen, 10977-015).

The donor plasmid was constructed by Gibson assembly using the following fragments: homology arms of ~1 kb on either side of the Cas9 cut site (two base pairs were deleted from the left arm immediately preceding the T2A maintain the open reading frame); a fragment containing *T2A-QF2-SV40* and *3xP3-dsRed*, PCR-amplified from a vector derived from *pBac-DsRed-ORCO_9kbProm-QF2* (gift of Chris Potter, Addgene #104877); a *pUC57* vector backbone digested with EcoRI and HindIII. Clones were sequenced verified and midiprepped using an endotoxin free midiprep kit (Machery-Nagel) and eluted in Ultrapure nuclease-free water (Invitrogen).

2000 *Ae. aegypti* embryos (LVP strain) were injected with a mixture of 300 ng/μL Cas9 protein (PNA Bio), 650 ng/μL dsDNA plasmid donor, and 40 ng/μL sgRNA. The progeny of 96 surviving G0 females were screened. Six potential founders were isolated, and one was verified to have a complete and in-frame insertion by PCR with the following primers: Forward 5’ GT GAGGGTGGTGT CGAATT AACT CTT 3’, Reverse 5’GTTAGGTCAGAGGTATCCCTGAACAT3’.

<u>*15xQUAS-mCD8-GFP*</u> was generated from an existing plasmid (a kind gift from Chris Potter, Addgene #104878). *Ae. aegypti* embryos (LVP strain) were injected with the plasmid and a PBac helper plasmid. Two independent lines were recovered.

<u>*15xQUAS-tdTomato-T2A-GCaMP6s*</u> was generated by Gibson assembly of the following PCR-amplified fragments: Plasmid backbone and Mos arms from *PUb-mCD8:GFP-T2A-dsRed:NLS-SV40* (described above); *15xQUAS from pBA C-ECFP-15xQUA S_TA TA-SV40* (Addgene #104875, gift of Chris Potter); *tdTomato-T2A-GCaMP6s* PCR-amplified from a vector synthesized (Genscript). *Ae. aegypti* embryos (LVP strain) were injected with the construct and a PBac helper plasmid. Two independent lines were recovered.

### Brain immunostaining

Dissection of adult brains and immunostaining was modified from previously used protocols^22,40^. 6-14 day-old mosquitoes were anesthetized on ice. Heads were carefully removed from the body by pinching at the neck with sharp forceps. Heads were placed in a 1.5 mL tube for fixation with 4% paraformaldehyde, 0.1 M Millonig’s Phosphate Buffer (pH 7.4), 0.25% Triton X-100, and nutated for 3 hr. Brains were then dissected out of the head capsule in ice cold Ca^+2^-,Mg^+2^-free phosphate buffered saline (PBS, Lonza, 17-517Q) and transferred to a 24-well plate. All subsequent steps were done on a low-speed orbital shaker. Brains were washed in PBS containing 0.25% Triton X-100 (PBT) at room temperature 6 times for 15 min. Brains were permeabilised with PBS, 4% Triton X-100, 2% Normal Goat Serum for ~48 hr (2 nights) at 4°C. Brains were rinsed once then washed with PBT at room temperature 6 times for 15 min. Primary antibodies were diluted in PBS, 0.25% Triton X-100, 2% Normal Goat Serum for ~48 hr (2 nights) at 4°C. The primary antibodies used in this experiment were anti-dmBrp (mouse; 1:50: NC82, DSHB) to label the synaptic neuropil and anti-GFP (Rabbit: 1:10,000; A11122, Life Technologies). Brains were rinsed once then washed in PBT at room temperature 6 times for 15 min. Secondary antibodies were diluted in PBS, 0.25% Triton X-100, 2% Normal Goat Serum for ~48 hr (2 nights) at 4°C. The primary antibodies used in this experiment were anti-mouse-Cy5 (1:250; Life Technologies A-10524) and anti-Rabbit-488 (1:500; Life Technologies A-11034). Brains were rinsed once then washed in PBT at room temperature 6 times for 15 min. Brains equilibrated overnight in Vectashield (Vector Laboratories H-1000), and were mounted in Vectashield.

### Ventral nerve cord immunostaining

6-14 day old mosquitoes were anesthetized on ice and the bodies were carefully removed from the heads by pinching at the neck with sharp forceps. The bodies were placed in a 1.5 mL tube for fixation with 4% paraformaldehyde, 0.1M Millonig’s Phosphate Buffer (pH 7.4), 0.25% Triton X-100, and nutated for 3 hr. Ventral nerve cords were dissected out of the body in ice cold PBS and transferred to a 24-well plate. All subsequent steps were the identical to the brain immunostaining protocol described above.

### ppk301>CD8:GFP immunostaining

*ppk301>CD8:GFP* expression was visualized in brains and ventral nerve cords using a Zeiss Inverted LSM 880 laser scanning confocal with a 25x / 0.8 NA immersion-corrected objective. Glycerol was used as the immersion medium to most closely match the refractive index of the mounting medium Vectashield. Brains were imaged at 2048 × 2048 pixel resolution in X and Y with 0.5 μm z-steps for a final voxel size of 0.1661 × 0.1661 × 0.5 μm. Ventral nerve cords were imaged at 1024 × 1946 pixel resolution in X and Y with 0.5 Lim z-steps for a final voxel size of 0.3321 × 0.3321 × 0.5 μm ventral nerve cord images were tiled and stitched with 10% overlap. Confocal images were processed in ImageJ (NIH).

3xP3 was used as a promoter to express fluorescent markers for transgene insertion, and care was taken to distinguish 3xP3 expression from the expression of the *ppk301-QF2* driver line. 3xP3 labels the optic lobes, as well as some cells in the dorsal brain. Figure 3j,k and Extended Data Figure 2b,d,f,h,j were cropped to remove 3xP3 expression. In some cases, we saw aberrant projections from the optic lobes that traversed the brain. These were sometimes seen in the driver alone and effector alone controls, both of which express 3xP3-ECFP, and we speculate that these are unrelated to *ppk301* expression. These patterns of 3xP3 are worth noting, but due to the distance from the subesophageal zone and ventral nerve cord, they do not have an impact on the results of this study. In some brains we observed faint unilateral projections from the subesophageal zone to the antennal lobe. We examined 8 female brains and 8 male brains for these projections and saw these projections in 5/8 brains. In males we did not see this projection in any of the 8 brains examined.

### Reference Brain 3D-Reconstruction

A reference *Ae. aegypti* brain from a 7-day old female LVP mosquito was fixed and immunostained for Brp as described above. It was imaged as described above except with 1 μm voxels and as a tiled scan with 10% overlap. 39 brains were imaged and the most complete and symmetric brain was chosen to serve as the template. Blind deconvolution was performed with AutoquantX3 software. The female brain was manually annotated using the segmentation and 3D reconstruction software ITK-SNAP. Regions were identified by the anatomical boundaries defined by Brp staining, and by homology to other insects, and named in accordance with revised insect brain nomenclature standards^23^. Structures without clear boundaries were excluded. A surface mesh of each region was exported into the data analysis and visualization software ParaView, which was used to generate the 3D reconstruction shown in Figure 3i.

### Mosquitobrains.org

The reference brain raw data and reconstruction are available on the website http://mosquitobrains.org. We have displayed this data as a brain atlas by creating an online ‘Brain Browser’ tool where users can scroll through the brain and highlight different regions. *Ae. aegypti* brains that are stained with the synaptic protein Brp can be warped onto this standard reference *Aedes aegypti* brain using the python code ClearMap^41^. To use ClearMap, all images must be acquired with square voxels. The standard brain was imaged with 1 μm voxel size. To use it as a template, all images must either be taken at this resolution, or down sampled to a final voxel size of 1 μm. Additional channels can be warped and registered onto the reference brain, provided that one channel is imaged as described above.

### T2A test construct validation

The previously characterized *Ae. aegypti* polyubiquitin promoter (Pub)^21^ was used to drive expression of both a membrane-bound variant of GFP (CD8:GFP) and a nuclear localization sequence fused to dsRED by separating these genes with the T2A ribosomal skipping element^19, 20^. This construct was expressed in a few cells in the adult brain, as well as in larval body-wall cells. We focused our analysis on the body-wall cells because their flat and compact shape made them amenable to examining the expression of our membrane and nuclear proteins.

Larvae were dissected by pinning the head and the tail to Sylgard plates (DowDupont) with insect pins, and making a long longitudinal cut along the dorsal surface of the body-all of the animal. The body wall was filleted open with four additional insect pins, and the organs were removed. The animals were fixed for 25 min in 4% paraformaldehyde, 0.1 M Millonig’s Phosphate Buffer (pH 7.4), 0.25% Triton X-100 at room temperature, and then rinsed 3 times in PBS. Larvae were transferred to a 1.5 mL tube and washed with PBT 6 times for 15 min while nutating at room temperature. Larvae were then incubated in Vectashield with DAPI overnight, and mounted in Vectashield for imaging. Larvae were imaged on a Zeiss Inverted LSM 880 laser scanning confocal with a 40X / 1.2NA oil immersion objective. The cells were imaged at a resolution of 2048 × 2048 pixels in a single confocal slice for a pixel size of 0.0692× 0.0692 μm. Images were processed in ImageJ (NIH).

### Visualization of ppk301-expressing cells in appendages

To visualize sensory neuron cell bodies in *ppk301-T2A-QF2, QUAS15xQUAS-tdTomato-T2A-GCaMP6* animals we dissected live sensory tissue, dipped in cold methanol for ~5 s, and mounted on a slide in glycerol. Appendages were viewed on a Zeiss Inverted lSm 880 laser scanning confocal with a 25x / 0.8 NA immersion corrected objective at a resolution of 1024 × 1024 pixels and a voxel size of 0.2076 μm × 0.2076 μm × 1μm.

### **Two-photon calcium imaging of female** ventral nerve cord

Calcium imaging was performed on an Ultima IV two-photon laser-scanning microscope (Bruker Nanosystems) equipped with galvanometers and illuminated by a Chameleon Ultra II Ti:Sapphire laser (Coherent). GaAsP photomultiplier tubes (Hamamatsu) were used to collect emitted fluorescence. Images were acquired with a 60X / 1.0N.A. Long Working Distance Water-Immersion Objective (Olympus) at a resolution of 256 × 256 pixels. Calcium imaging experiments were performed on female mosquitoes that were 7-11 days post-eclosion. Mosquitoes were fed a human-blood-meal 96-108 hours prior to imaging and were not giving access to an egg laying substrate so that they were gravid at the time of imaging. Gravid females were anesthetized at 4°C for dissection. The wings were removed and the mosquito was fixed to a custom Delrin plastic holder with UV-curable glue (Bondic). The mosquito was inserted into a hole in the holder, such that the ventral thorax, including all coxae, were exposed above the surface of the holder, with the rest of the mosquito below. The mosquito was secured with a few points of glue (Bondic) on the abdomen, thorax and head. The leg that was presented with water remained free of glue to prevent damage to the tissue. Once the mosquito was secured to the plate, one of the forelegs was inserted into a small diameter tube that was secured to the bottom of the plate and could later be attached to the fluidics apparatus used for stimulus delivery (described below). The top of the dish was then filled with external saline. The recipe we used is based *Drosophila melanogaster* imaging saline:103 mM NaCl, 3 mM KCl, 5 mM 2-[Tris(hydroxymethyl)methyl]-2-aminoethanesulfonic acid (TES), 1.5 mM CaCl_2_, 4 mM MgCl_2_, 26 mM NaHCO_3_, 1 mM NaH_2_PO_4_, 10 mM trehalose, 10 mM glucose, pH 7.3, osmolality adjusted to 275 mOsm/kg). The coxae were gently spread from the midline and secured in dental wax. The cuticle was removed above the prothoracic ganglia using very sharp forceps. Opaque non-neural tissue, primarily fat cells and muscle, was removed if they obstructed the ventral nerve cord. Great care was taken not to damage the *ppk301*-expressing nerves running from the legs into the ventral nerve cord. These run up the posterior-ventral region of the leg and are extremely superficial. tdTomato fluorescence was examined before imaging to verify that the nerves were intact. If an animal did not respond to water, 200 mOsm/kg, or 300 mOsm/kg NaCl it was discarded.

The preparation was secured to the stage using a custom laser cut acrylic holder. A single plane through the centre of the prothoracic neuropil was scanned at 4.22 fps with a 920 nm excitation wavelength imaged through a 680 nm shortpass IR blocking filter, a 565 nm longpass dichroic and 595/50 nm or 525/70 nm bandpass filters. GCaMP6s and tdTomato emission was collected simultaneously for 70 frames per trial. Each concentration was delivered at least 3 times per animal, and each animal was exposed to 0, 200, and 300 mOsm/kg NaCl. Imaging remained stable during the duration of the imaging session in all animals that were included in this study. We did not notice a decrease in the response to stimuli over time. Before beginning the experiments using multiple concentrations of salt, we imaged animals with repeated water delivery, and saw no desensitization to the response to water over 10 presentations (data not shown).

### Liquid Delivery

Liquids were delivered to a single foreleg of the mosquito using a custom built low-volume fluidics device. Piezoelectric diaphragm micropumps and their controller (Servoflo) were run by an Arduino using a code written for this purpose. A custom manifold was milled for small volume liquid delivery. The mosquito was illuminated with an IR light and liquid delivery was monitored using an IR camera. The liquid coated the tarsal and tibial segments and was retracted immediately after imaging each sweep. We waited at least 1 min between each trial. We only observed responses in the prothoracic region ipsilateral to the stimulus delivery.

### Data Analysis

All image processing was done using FIJI/ImageJ (NIH). Further processing was done using Excel and Prism (GraphPad). Regions of interest were selected based on the tdTomato fluorescence intensity, and used for analysis of GCaMP6s signal. All traces with motion, as determined by tdTomato fluorescence instability, were discarded. A Gaussian blur with a sigma value of 1 was performed on the GCaMP6s signal. In the calculation of ΔF/F, 6 frames were averaged before stimulus presentation to determine the baseline fluorescence. To determine F_max_, the average of 3 frames at the peak after stimulus delivery was determined for each sweep. The one-way ANOVA with Tukey’s HSD presented in Figure 4e compared the average values from a single animal for each concentration.

### Statistical analysis

All statistical analyses were performed using Prism (GraphPad) or R version 3.5.0 ^37^. Data collected as percentage of total are shown as median with interquartile range and data collected as raw value are shown as mean ± SEM or mean ± SD. Details of statistical methods are reported in the figure legends.

### Data Availability Statement

All plotted data (with the exception of raw video files) are available in the Supplemental Data File, and behaviour assay schematics and egg-laying counting image analysis scripts can be found at https://github.com/VosshallLab/MatthewsYoungerVosshall2018. Plasmids are available from Addgene.

---

**Extended Data Figure 1.**
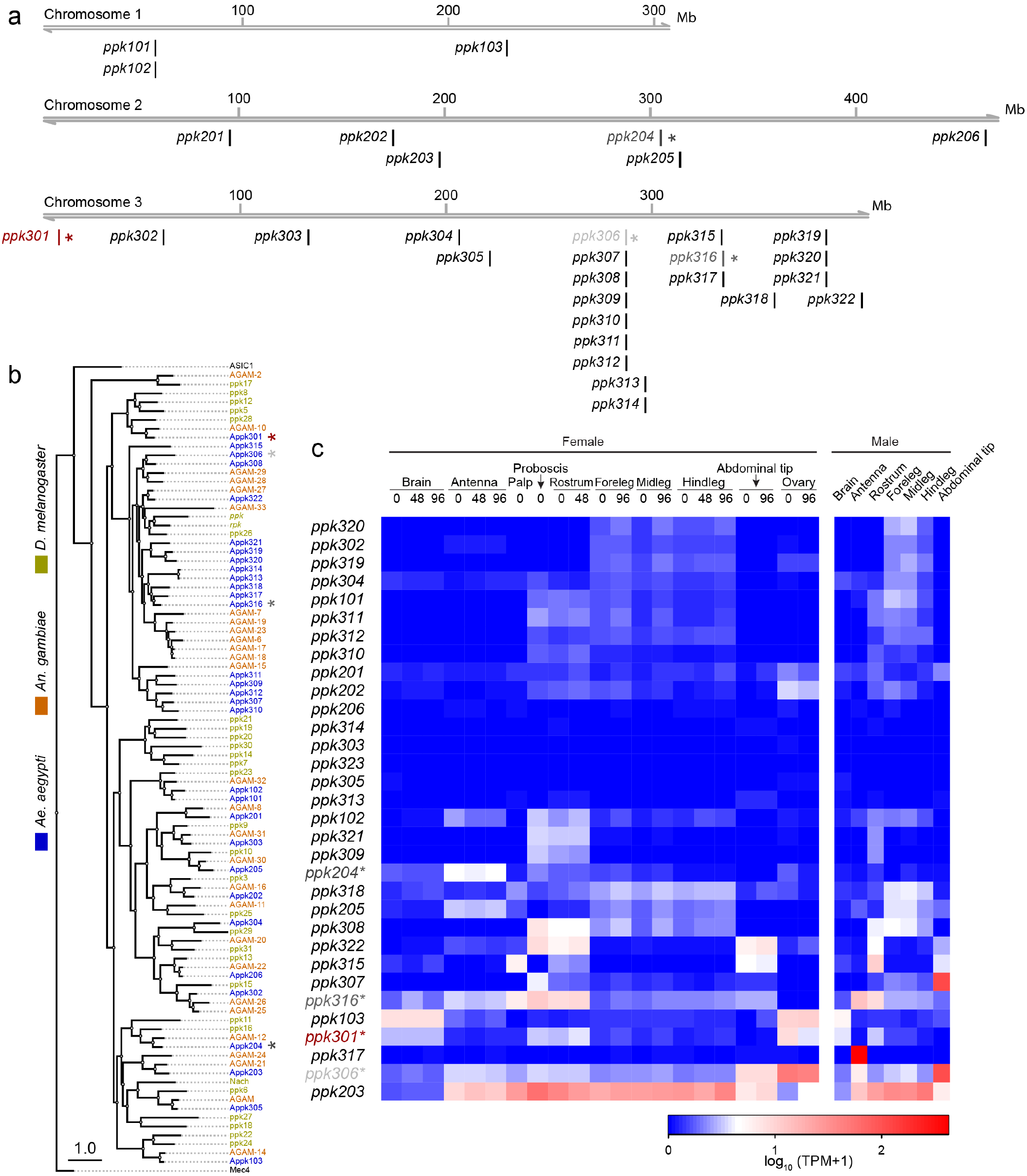
Genomic organization and tissue-specific expression of *Ae. aegypti ppk* ion channels. **a,** Genomic organization of ppk ion channels on the three chromosomes of *Ae. aegypti.* **b,** Phylogenetic tree of ppk ion channel proteins from *Ae. aegypti, Anopheles gambiae*, and *Drosophila melanogaster*. Scale bar = 1.0 substitution/site. **c,** Gene expression of *ppk* ion channels across tissues. Data originally generated in a survey of gene expression across neural tissues^34^ were aligned to the AaegL5 genome^35^. Values represent mean of multiple replicates for each tissue.

**Extended Data Figure 2.**
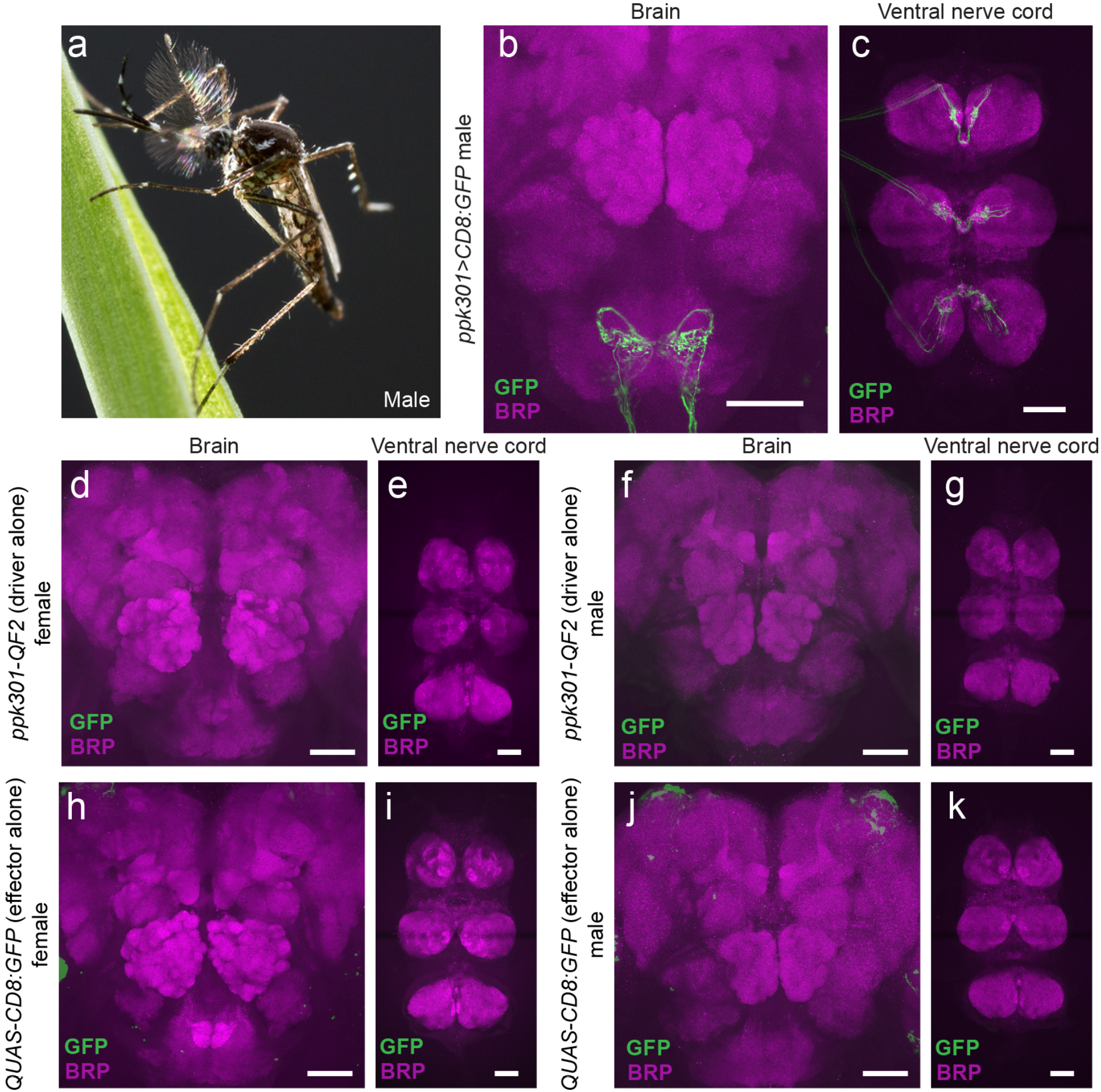
Male expression of *ppk301>CD8:GFP* and QF2/QUAS controls for *ppk301-QF2* reagent. **a,** Male *Ae. aegypti* mosquito. Photo: Alex Wild. **b, c,** Expression of *ppk301>CD8:GFP* in male brain (**b**) or ventral nerve cord (**c**). **d, e, f, g,** Female brain (**d**) or ventral nerve cord (**e**) and male brain (**f**) or ventral nerve cord (**g**) in driver-only *ppk301-QF2* control animals. **h, i, j, k,** Female brain (**h**) or ventral nerve cord (**i**) and male brain (**j**) or ventral nerve cord (**k**) in responder-only *QUAS-CD8:GFP* control animals. In all panels, Brp is in magenta and CD8:GFP in green. Scale bars: 50 μm.

